# Novel reference transcriptomes for the sponges *Carteriospongia foliascens* and *Cliona orientalis* and associated algal symbiont *Gerakladium endoclionum*

**DOI:** 10.1101/2020.06.26.156463

**Authors:** Brian W. Strehlow, Mari-Carmen Pineda, Carly D. Kenkel, Patrick Laffy, Alan Duckworth, Michael Renton, Peta L. Clode, Nicole S Webster

## Abstract

Transcriptomes from sponges are important resources for studying the stress responses of these ecologically important filter feeders, the interactions between sponges and their symbionts, and the evolutionary history of metazoans. Here, we generated reference transcriptomes for two common and cosmopolitan Indo-Pacific sponge species: *Carteriospongia foliascens* and *Cliona orientalis.* We also created a reference transcriptome for the primary symbiont of *C. orientalis* – *Gerakladium endoclionum*. To ensure a full repertoire of transcripts were included, clones of each sponge species were exposed to a range of individual stressors: decreased salinity, elevated temperature, elevated suspended sediment concentrations, sediment deposition and light attenuation. RNA extracted from all treatments was pooled for each species, using equal concentrations from each clone. Sequencing of pooled RNA yielded 409 and 418 million raw reads for *C. foliascens* and *C. orientalis* holobionts (host and symbionts), respectively. Reads underwent quality trimming before assembly with Trinity. Assemblies were filtered into sponge-specific or, for *G. endoclionum*, symbiont-specific assemblies. Assemblies for *C. foliascens, C. orientalis*, and *G. endoclionum* contained 67,304, 82,895, and 28,670 contigs, respectively. Contigs represented 15,248-37,344 isogroups (∼genes) per assembly, and N50s ranged 1,672-4,355 bp. Gene ortholog analysis verified a high level of completeness and quality for sponge-specific transcriptomes, with an average 93% of core EuKaryotic Orthologous Groups (KOGs) and 98% of single-copy metazoan core gene orthologs identified. The *G. endoclionum* assembly was partial with only 56% of core KOGs and 32% of single-copy eukaryotic core gene orthologs identified. These reference transcriptomes are a valuable resource for future molecular research aimed at assessing sponge stress responses.

## Data Description

Sponges, phylum Porifera, represent one of the oldest lineages of multicellular animals [1], hence investigating the transcriptomes of different sponge species can provide insight into the evolution of metazoans and their gene expression profiles. Furthermore, sponges have an uncertain future in the face of global climate change [2,3] as well as local stressors including coastal development, altered hydrological processes, and increased runoff of nutrients, pesticides and sediments [4–7]. Transcriptomic analysis of sponges that have been exposed to different environmental conditions would improve our understanding of the sponge molecular stress response pathways and enhance our ability to effectively manage these ecologically important filter feeders. Although there are approximately 9,000 described sponge species [8], to date only ∼35 species have published transcriptomes [9-25] and only ∼10 have published genomes [10,16,26–29].

In this study, we assembled the transcriptomes of two common and widely distributed Indo-pacific sponge species – *Carteriospongia foliascens* and *Cliona orientalis*. Both are emerging model species that have been extensively used to study the physiological and ecological effects of environmental stressors on sponges [30–37]. *C. foliascens* and *C. orientalis* are only the second members of their respective orders (Dictyoceratida and Clionaida) to have a reference transcriptome sequenced. Whilst both *C. foliascens* and *C. orientalis* host diverse populations of bacterial symbionts, e.g. [32], *C. orientalis* additionally hosts an abundant population of eukaryotic Symbiodiniaceae, *Gerakladium endoclionum* [38,39], which comprises up to 96% of its algal symbiont community [37]. We used sequences generated from the *C. orientalis* holobiont, i.e. host and symbiont, to construct a partial reference transcriptome for *Gerakladium endoclionum*. Matching host and symbiont transcriptomes provide a valuable tool to understand the holobiont response to changing environmental conditions and determine the cause-effect pathways for declining host health with environmental change. These data contribute substantially to available poriferan genetic resources and advance the development of these two sponge species as model systems for field and laboratory studies.

## Methods

### Samples and sequencing

Samples of *C. foliascens* and *C. orientalis* were collected in May 2015 from Fantome Is. (S 18°41.028⍰ E 146° 30.706) and Pelorus Is. (S 18°32.903’ E 146° 29.172’), respectively, in the central Great Barrier Reef under permits G12/35236.1 and G13/35758.1. As *C. orientalis* is a bioeroding sponge that encrusts and erodes coral skeletons, five *C. orientalis* cores (∼5 cm in diameter) were collected using an air-drill from a single individual, i.e. cloned, growing on a dead colony of *Porites* sp. An individual of *C. foliascens* was cut (cloned) into five pieces as in [32]. Sponges were healed and acclimated under natural light and flow-through seawater for 4 weeks before experiments were performed.

In order to capture the full complement of gene expression within the reference transcriptomes, sponges were subjected to five different treatments at the Australian Institute of Marine Science (AIMS) National Sea Simulator: i) decreased salinity, ii) elevated temperature, iii) elevated suspended sediment concentrations (SSCs) and sediment deposition, iv) light attenuation and v) no stress control. Sponge clones were used across all treatments to control for genotype, i.e. one genotype was used per species. Two clones of each species were used for each treatment. In the salinity stress treatment, salinity was decreased from 35 to 22 parts per thousand (ppt) by gradually adding flow-through reverse osmosis (RO) water to the system. Salinity was held constant at 22 ppt for 2 d with flow-through seawater maintained at 600 mL min^-1^. In the heat stress treatment, sponges were exposed to a constant temperature of 32.5°C for 1 d using methods described in [32]. In the sediment treatment, sponges were exposed to elevated SSCs at 200 mg L^-1^ for 1 d as in [40,41], using sediments described therein. In the deposition experiment, sedimentation was approximately 40 mg cm^-2^, measured using SedPods [42] and sponges were left covered with sediment for 1 d. In the light attenuation treatment, sponges were kept in complete darkness for 2 d. Immediately after each treatment, a sample of sponge tissue (∼1 cm^3^) was flash frozen in liquid nitrogen and stored at -80°C [43] for RNA extraction and sequencing (RNA-seq). After exposure to the decreased salinity and darkness treatments, *Cliona orientalis* was visibly bleached after 2 d, but *C. foliascens* did not exhibit any colour change. Sponges were not visibly affected, e.g. no bleaching or necrosis, by the sediment exposure or elevated seawater temperature.

Approximately 50 mg of each sponge clone was excised and ground using a mortar and pestle. Grinding was performed under a thin layer of liquid nitrogen to limit RNA degradation. All tools were rinsed in ethanol followed by RNase Zap (Sigma-Aldrich, USA) to remove contamination and deactivate RNA degrading enzymes. Total RNA was isolated using the Zymo ZR RNA miniprep kit (Zymo Research, USA), with in-column DNAse digestion, according to the manufacturer’s protocol. Total RNA was subsequently cleaned using the Zymo RNA Clean and Concentrator kit (Zymo Research, USA), following the Manufacturer protocol.

Total RNA quality was checked using gel electrophoresis and spectroscopy (NanoDrop 2000c Spectrophotometer, Thermo Fisher Scientific, USA), and quantified using a Quant-iT Ribogreen Assay (Thermo Fisher Scientific, USA). For each sponge species, the RNA from individual treatments was combined in equal amounts (740 ng for *C. foliascens* and 1,440 ng for *C. orientalis*) from all sponge clones, to a final total RNA concentration of 3.7 µg in 40 µl of Dnase and Rnase free water (93 ng µl^-1^) for *C. foliascens* and 7.2 µg in 55 µl of Dnase and Rnase free water (160 ng µl^-1^) for *C. orientalis*. For *C. foliascens* and *C. orientalis* respectively, RNA had a ratio of absorbance at 260 nm to 280 nm (A260/A280) of 1.88 and 2.02 and an A260/A230 ratio of 1.11 and 1.67. To isolate eukaryotic messenger RNA (mRNA), a TruSeq Stranded mRNA-seq sample prep was performed prior to sequencing. The mRNA was sequenced across two lanes of Illumina HiSeq2500 at the Ramaciotti Centre for Genomics (University of New South Wales, Sydney, Australia) to generate 2 x 100 base pair (bp) paired-end (PE) rapid runs.

### Transcriptome assembly and annotation

Sequencing produced 409 and 418 million raw reads for *C. foliascens* and *C. orientalis*, respectively (Table 1). Reads were trimmed and assembled using publicly available scripts [44,45] and the protocol detailed in [46]. Briefly, reads < 50 bp long were removed along with reads containing a homopolymer run of adenine (A) longer than 9 bases using *fastx_toolkit* [47], and only reads with a PHRED quality score >20 over 80% of the read were retained. TruSeq sequencing adapters and PCR duplicates were also removed [44]. The remaining filtered, high quality reads (32.5, 50.5 million paired reads and 2.9, 8.7 million unpaired reads for *C. foliascens* and *C. orientalis*, respectively) underwent *de novo* assembly into contigs using Trinity v 2.8.5 [48]. Data processing and assembly was performed at AIMS in Townsville using a high-performance computer (HPC) and on ABACUS 2.0 at the Danish e-Infrastructure Cooperation (DeiC) National HPC Center.

**Table 1.** Assembly statistics for the *de novo* transcriptomes.

|  | *Carteriospongia foliascens*<br>holobiont | *Cliona orientalis*<br>holobiont |  |
| --- | --- | --- | --- |
| N raw reads (x10<sup>6</sup>) | 409 | 418 |  |
| N qual filtered: PE, SE (x10<sup>6</sup>) | 32.5, 2.9 | 50.5, 8.7 |  |
| N contigs holobiont | 146,510 | 225,126 |  |
|  | *Carteriospongia foliascens* | *Cliona orientalis* | *Gerakladium endoclionum* |
| N contigs target species only | 67,304 | 82,895 | 28,670 |
| Mean GC content target species only | 40.2 | 45.5 | 59 |
| N genes | 15,248 | 37,344 | 21,566 |
| Mean contig length (bp) | 3,024 | 1,756 | 1,375 |
| N50 (bp) | 4,355 | 2,369 | 1,672 |
| % Annotated | 64 | 77 | 39 |
| % core KOGs | 97.9 | 98.7 | 56 |
| BUSCOs |  |  |  |
| N complete (%) | 908 (92.8) | 921 (94.2) | 98 (32.3) |
| N partial (%) | 14 (1.48) | 16 (1.94) | 16 (5.28) |
| N missing (%) | 56 (5.73) | 38 (3.89) | 189 (62.8) |

Following assembly, additional quality control was performed to ensure that only target transcripts, i.e. derived from *C. foliascens, C. orientalis* or *G. endoclionum*, were included in their respective reference transcriptomes [13,46]. First, contigs less than 400 bp were removed and ribosomal RNA (rRNA), mitochondrial RNA (mtRNA), Symbiodiniaceae, and other non-metazoan (e.g. bacteria) sequences were identified using a series of hierarchical BLAST [49] searches. Transcriptomes were further blasted (BLASTn) against the *A. queenslandica* rRNA database (SILVA: ACUQ01015651) [50], which was the most complete Poriferan rRNA database. Contigs with a bit-score >45 were removed, i.e. 9 and 10 sequences in *C. foliascens* and *C. orientalis*, respectively. This process was repeated using the *A. queenslandica* mitochondrial genome (NCBI: NC_008944.1 REF), resulting in 61 and 27 sequences being removed from the *C. foliascens* and *C. orientalis* assemblies respectively.

Remaining contigs were blasted (BLASTx) against the most complete Poriferan (*A. queenslandica*, aqu2.1_Genes_proteins.fasta) [28,51] and *Symbiodinium kawagutii* (Symbiodinium_kawagutii.0819.final.gene.pep) [52] predicted proteomes and the NCBI nonredundant (nr) protein database (downloaded September 2019). In order to be included in a sponge-specific assembly, contigs had to return a more significant match (E value ≤ 10^−5^) to the *A. queenslandica* proteome compared to blast results from the *S. kawagutii* proteome *and also* match a metazoan sequence in the nr database or have no match in the nr database, as described in [13]. Sequences with no match to either proteome were excluded from the final sponge assemblies [13], a stricter exclusion procedure than used in prior invertebrate transcriptome assemblies [46]. For the *C. orientalis* holobiont, sequences matching the *S. kawagutii* proteome more closely than the *A. queenslandica* proteome (E value ≤ 10^−5^) and matching to the phylum chromerida in the nr database (or having no match in the nr database) were included in the final *G. endoclionum* assembly. Although *C. foliascens* does not contain intracellular Symbiodiniaceae, the decontamination step was also performed in order to remove any potential algal contamination in the sample, resulting in only a few (1,520, 1% of total number of contigs) contaminating sequences being removed.

Within each of the three transcriptomes, contigs were assigned to isogroups (∼genes) and given gene names and gene ontologies (GO) [53] following the protocol previously described in [44,54]. Briefly, the transcriptomes underwent BLAST pairwise sequence comparison (BLASTx) to the UniProt Knowledgebase (UniprotKB/Swiss-Prot) database [55]. Significant BLASTx results (E value ≤ 10^−4^) were used by CDS_extractor_v2.pl [56] to extract and identify protein coding sequences. Functional annotations were assigned to isogroups based on orthologous comparisons to the eggNOG 4.5 database [57] using eggnog-mapper [58]. Kyoto Encyclopedia of Genes and Genomes (KEGG) ids were also assigned to isogroups using the KEGG Automatic Annotation server (KAAS) [59]. The guanine-cytosine (GC) content of transcriptomes was calculated using the BBMap package (Joint Genome Institute, USA) [60]. Transcriptome completeness was assessed by Benchmarking Universal Single-Copy Orthologs (BUSCO) analysis [61] on gVolante [62].

### Assembly evaluation and quality control

The holobiont assemblies of *C. foliascens* and *C. orientalis* contained 225,126 (N50 = 1,284) and 146,510 (N50 = 1,949) contigs greater than 400bp in length (Table 1). After data partitioning, 67,304 and 82,895 contigs, for *C. foliascens* and *C. orientalis* respectively, were considered the ‘sponge-specific’ transcriptome assemblies. The partitioned *G. endoclionum* transcriptome isolated from the *C. orientalis* holobiont comprised 28,670 contigs (Table 1). The *C. foliascens, C. orientalis*, and *G. endoclionum* transcriptomes contained 15,248, 37,344, and 21,566 isogroups, respectively, with mean lengths of 3,024 (N50 = 4,355), 1,756 (N50 = 2,369), and 1,375 (N50 = 1,672) bp (Table 1). The number of isogroups identified in the *C. foliascens* and *C. orientalis* transcriptomes was comparable to previously published sponge transcriptomes which have reported ∼11,000-60,000 expressed genes [15,23,63], although there is considerable variation across species. The *G. endoclionum* transcriptome, containing 28,670 isogroups was comparable in size to the previously published *S. kawagutii* genome (36,850 genes) [52] and previously published Symbiodiniaceae transcriptomes, ranging in size from 23,777-26,986 expressed genes [64]. The respective GC content of each assembly was 40.2, 45.5, and 59%, matching reported values for metazoans (35-55% [17,27,65]) and Symbiodiniaceae (45-65% [65]). For *C. foliascens* and *C. orientalis*, the percentage of genes assigned a name or GO terms was 64 and 77%, respectively (Table 1), also comparable to other sponge transcriptomes (30-70% [15]) and those of other non-model metazoans (25-62% [14,46]). In comparison to the annotated sponge transcriptomes, only 39% of *G. endoclionum* isogroups could be assigned function or GO term annotations, however this is consistent with functional annotation of other intracellular Symbiodiniaceae transcriptomes, where between 34-44% of genes were assigned GO terms [64]. The isogroups for *C. foliascens, C. orientalis*, and *G. endoclionum* were assigned 3,641, 5,339 and 2,191 unique KEGG annotations respectively.

The representative transcriptomes for *C. foliascens* and *C. orientalis* are considered largely complete based on BUSCO analysis (92.8% and 94.2% complete, respectively) and the representation of nearly all core eukaryotic Orthologous Groups (KOGs) (97.9% and 98.7% respectively) (Table 1). BUSCO analysis of the transcriptome of *G. endoclionum* was 32.3% complete and 56% of core KOGs were identified (Table 1). A reduced BUSCO completeness in transcriptomes isolated from intracellular Symbiodiniaceae in corals (33-42%) has been previously reported [66]. The current *G. endoclionum* transcriptome contained 86% more isogroups than the Symbiodiniaceae transcripts identified within the transcriptome assembly of the closely related sponge holobiont, *Cliona varians* [17]. *C. varians* also hosts a congeneric intracellular photosymbiont, *Gerakladium spongiolum* [38]. Therefore, the current transcriptome for *G. endoclionum* was considered useful for future studies, at least for conditions *in hospite*.

### Re-use potential

These reference transcriptomes were assembled to facilitate sponge holobiont research aimed at exploring how both host and symbionts respond to changing environmental conditions. The transcriptomes can be used for studies involving Tag-based RNAseq (TagSeq) [67], a highly accurate [68] and cost-effective sequencing technique for large sample sets. Output files are also formatted for Rank-based Gene Ontology analysis of gene expression data (GO_MWU, [69]), and for Functional Summary and Meta-Analysis of Gene Expression Data (KOGMWU, [70]).

### Availability of supporting data

All data, including raw reads, can be accessed here: https://www.dropbox.com/sh/82ue5l16n4xzxww/AABENUi-Cdbm_z-6X4Gj3qICa?dl=0. Raw data has also been deposited on NCBI’s SRA under accession numbers PRJNA639714 and PRJNA639798 for the *C. orientalis* holobiont and *C. foliascens*, respectively.

## List of abbreviations

BUSCO: Benchmarking Universal Single-Copy Orthologs;
KEGG: Kyoto Encyclopedia of Genes and Genomes;
KOG: EuKaryotic Orthologous Groups;
TagSeq: Tag-based RNAseq.

## Declarations

### Ethics notes and consent for publication

Not applicable.

### Authors’ contributions

BWS, MCP, CDK, AD, MR, PC, and NSW conceived and designed the experiments; BWS, MCP, and CDK performed the experiments and laboratory work; BWS performed bioinformatics analyses with extensive help and training from CDK and PL. BWS wrote the first draft. BWS, MCP, CDK, PL, AD, MR, PC, and NSW contributed to revisions of various drafts and approved the final manuscript.

## Acknowledgements

The authors are grateful to E. Botté and G. Millar for their assistance in technical aspects related to computing. They also thank the staff at the AIMS National Sea Simulator for their expertise and assistance in the tank-based experiments.

